# Non-selective cation permeation in an AMPA-type glutamate receptor

**DOI:** 10.1101/2020.06.21.162735

**Authors:** Johann Biedermann, Sebastian Braunbeck, Andrew J. R. Plested, Han Sun

## Abstract

Fast excitatory synaptic transmission in the central nervous system relies on the AMPA-type glutamate receptor (AMPAR). This receptor incorporates a non-selective cation channel which is opened by the binding of glutamate. Although the open pore structure has recently became available from cryo-electron microscopy (Cryo-EM), the molecular mechanisms governing cation permeability in AMPA receptors are not understood. Here, we combined microsecond molecular dynamics (MD) simulations on a putative open state structure of GluA2 with electrophysiology on cloned channels to elucidate ion permeation mechanisms. Na^+^, K^+^ and Cs^+^ permeated at physiological rates, consistent with a structure that represents a true open state. A single major ion binding site for Na^+^ and K^+^ in the pore represents the simplest selectivity filter (SF) structure for any tetrameric cation channel of known structure. The minimal SF comprised only Q586 and Q587, and other residues on the cytoplasmic side formed a cone-shaped void that lacked major interactions with ions. We observed Cl^-^ invasion of the upper pore, explaining anion permeation in the edited form of GluA2. A permissive architecture of the SF accommodated different alkali metals in distinct solvation states to allow rapid, non-selective cation permeation, and co-permeation by water. Simulations suggested Cs^+^ uses two equally populated ion binding sites in the filter and we confirmed with electrophysiology of GluA2 that Cs^+^ is more permeant than Na^+^, consistent with serial binding sites preferentially driving selectivity.

**Significance Statement:** AMPA-type glutamate receptors (AMPARs) are key actors in neurotransmission, making the final step in a relay of excitability from one brain cell to another. The receptor contains an integral ion channel, which, when opened by neurotransmitter binding, permits sodium and other cations to cross the cell membrane. We investigated permeation of sodium, potassium and caesium in an AMPAR at the atomistic level using a computational molecular dynamics approach on a structure with the ion channel pore in a presumably open state. We determined that the region selecting between cations is the simplest of any channel of this type. Distinct from ion channels that select single ion species, cations are never fully dehydrated and have only one major ion binding site in the filter. Simulations suggested two similar binding sites for caesium, and studies of AMPARs in mammalian cell membranes showed that this makes caesium more permeant than sodium.

## Introduction

Glutamate receptor ion channels are found at synapses throughout the vertebrate nervous system, where they convert sub-millisecond glutamate signals into cation currents. Advances in structural biology have provided molecular scale maps of their ion pores, permitting comparison with a burgeoning menagerie of structures from related ion channels. It has been comparatively difficult to obtain candidate open pore structures of glutamate receptors, with the notable exceptions being from single particle cryo-electron microscopy (Cryo-EM) of complexes between GluA2 and Stargazin (1, 2). However, it is unclear from simple inspection of these structures whether a) the ion pore is conductive, or b) it is open to its fullest extent, or to the highest conductance level. However, a substantial body of electrophysiological work provides good benchmarks for how these structures should behave. For example, canonical measurements of channel conductance and reversal potentials show that alkali earth cations from sodium (Na^+^) up to caesium (Cs^+^) should permeate GluA1 (3) and unedited GluA2 (4) approximately equally well and the single channel conductance of the full level of GluA2 (Q) should be considerable (∼30 pS, ref. 5).

Most computational work on ion permeation through channels has been focused on simple, selective potassium (K^+^) channels like KcsA (6, 7), being the first reported crystal structures of ion channels (8). Thanks to their minimal sequences, these channels demand little computational overhead. Their key structural features are two membrane spanning helices and a reentrant loop forming a narrow selectivity filter (SF) for permeant ions. This core motif defines a superfamily of tetrameric and pseudotetrameric channels that encompass selective, semi-selective and non-selective cation conductances. In common with many eukaryotic channels, the AMPA type glutamate receptor has a pore domain whose gating state is controlled by substantially larger domains outside the membrane (amino terminal domain, ATD and ligand binding domain, LBD, Figure 1A) which account for about 75% of the protein mass. The large size presents a challenge for conventional molecular dynamics (MD) simulations, with the AMPA receptor being about 6 times bigger than KcsA.

**Figure 1.**
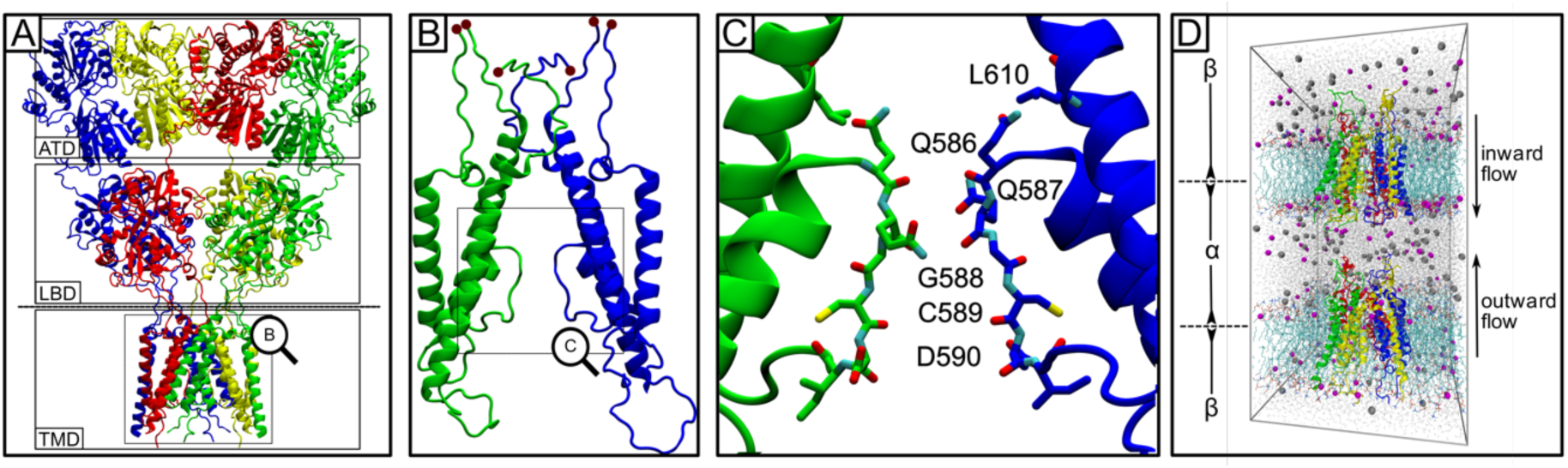
AMPA receptor and the simulation setup. (A) The activated open state of the AMPA receptor from Cryo-EM (PDB ID: 5WEO) (1) with Stargazin molecules removed. The receptor is composed of amino terminal, ligand binding and transmembrane domains (ATD, LBD and TMD). (B) The TMD and linkers to the LBD layer were included the MD simulations. The sites where the linkers were truncated and physically restrained (see Methods) are marked with red balls. Two out of the four subunits are shown. (C) The selectivity filter region of the AMPA receptor pore, with key residues labelled. Again, only two diagonally-opposed subunits are drawn. (D) The computational electrophysiology setup was composed of two tetrameric AMPA channels, each embedded in a separate POPC lipid bilayer, solvated with water molecules and ions. A small cation imbalance between the two compartments *α* and *β* was maintained during the simulations. The resulting gradient gave a transmembrane potential to drive ion permeation.

It remains to be seen to what extent key features of ion permeation elucidated in prokaryotic channels (selectivity, discrete sites, desolvated ions and block by divalent ions) are widely applicable in channels with more substantial architectures. Recent experimental and MD work on non-selective prokaryotic channels like NaK and NaK-CNG suggests that ion permeation in these channels differs substantially from classical K^+^ selective ion permeation (9–11). The SF of non-selective cation channels is much more flexible, with fewer ion binding sites leading to distinct conduction mechanisms and hydration states for Na^+^ and K^+^ when passing through the filter. In the AMPA receptor, in one activated open Cryo-EM structure of GluA2 (1), density for a presumptive hydrated sodium ion was observed adjacent to the unedited Q586 residue. In the structure of edited GluA2 (Q586R) with Stargazin (2), ions were absent. The closed state structure of a GluA1:A2 heteromer featured strong density of unknown identity adjacent to C589 (12). Whether the paucity and heterogeneity of resolved ions is due to a lack of order in the filter region, the lack of detail in the coulomb potential density or a true deficit of ions remains unclear. However, observations in these AMPA receptor structures are in marked contrast to the Cryo-EM structures of several other non-selective cation channels such as hyperpolarization-activated cyclic nucleotide-gated (HCN) (13) and cyclic nucleotide-gated (CNG) channels (14) where two or three bound ions were visible. Further context comes from canonical K^+^ -selective channels which feature up to 4 ions in a row (15) and crystal structures of Ca^2+^ -selective TRP channels where two Ca^2+^ ions were readily resolved (16).

Here we used MD-based computational electrophysiology to examine ion permeation through the mammalian AMPA receptor ion channel. We determined that the published structure is stably open, suitably detailed for MD simulations and probably represent a native fully-open state. We identified a minimal selectivity filter consisting of a single ion binding site that does not dehydrate ions. In simulations, Cs^+^ co-opted a secondary binding site. Consistent with multiple sites promoting ion selectivity, electrophysiology of AMPARs in HEK cells showed that Cs^+^ is more permeant than Na^+^.

## Results

### Monovalent cation permeation

Our starting structure for MD simulations was the AMPAR channel in its activated conformation (PDB ID: 5WEO) (1), which we embedded into a palmitoyloleoyl phosphatidylcholine (POPC) lipid bilayer (Figure 1D). To reduce the number of atoms in the simulations, we removed the extracellular ATD and LBD and the four Stargazin molecules decorating the periphery of the channel. To retain the open channel conformation, we held the transmembrane domain (TMD) of AMPAR open by physically restraining the end of the truncated linkers (Figure 1B, see Methods for details). This approach, while facile, was highly reproducible, and the channel did not close spontaneously during any simulation run (Figure 2A). Simulations of ion conduction were performed with the computational electrophysiology method (17, 18). In this setup, the simulation box contains two membranes that define two compartments and a voltage difference across each membrane is created by an ion gradient that is maintained alchemically during the simulations (Figure 1D).

**Figure 2.**
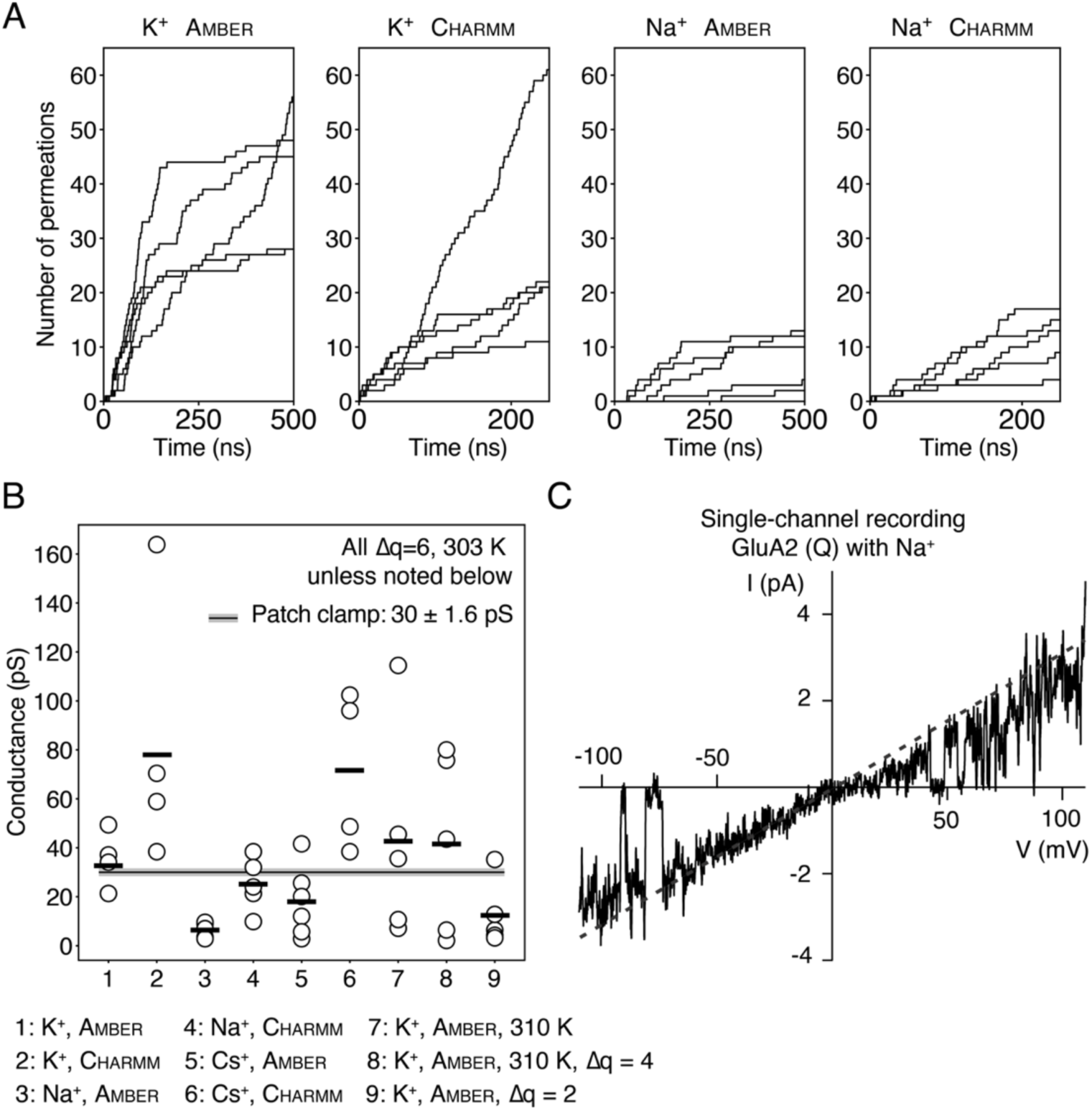
Ion conductances in GluA2. (A) Cumulative ion permeation events in the GluA2 pore for individual trajectories of K^+^ and Na^+^ simulations with the respective force fields. The simulations were performed at 303 K using an ion imbalance of 6e^−^ between two compartments *α* and *β*. (B) Conductances derived from the K^+^, Na^+^ and Cs^+^ simulations performed with AMBER99sb and CHARMM36 force fields. Simulations were carried out at different temperature and with different ion imbalance between two compartments *α* and *β*. The experimental value derived from single-channel recordings for Na^+^ is displayed as a grey line. (C) Outside-out patch clamp electrophysiology of a single GluA2 (Q) channel in symmetrical Na^+^. The voltage ramp was made in the presence of 10 mM glutamate and 100 *µ*M CTZ. The dashed line shows the fitted chord conductance, which for this patch was 33 pS.

Classical electrophysiology experiments suggest AMPARs are broadly non-selective over different alkali metal cations, although most measurements were not made on GluA2 (3, 19). We began by performing a series of ion permeation simulations with the principal biological ion species, Na^+^ and K^+^. To ensure the robustness of our observations, we undertook simulations at different temperatures, with different force fields and different driving forces on the ions (see Discussion and Methods). The conductances of the AMPA channel pore were directly calculated from the ion permeations observed in these simulations. We counted cations that traversed the entire narrow region of the pore domain delineated by the ^586^QQGCDI^591^ sequence, summarized in Table 1 and Figure 2B. Outside of this region, ion movements lacked order and made only rare interactions with the channel leading to a largely flat energy landscape (Figure S1). In all trajectories, the channel pore remained conductive for Na^+^ or K^+^ during the entire 500 ns run, as exemplified by the ion track plots in Figure 2A. In a typical 500 ns trajectory with a transmembrane potential of 450 mV, we recorded over 40 K^+^ permeation events, corresponding to a conductance of 32 pS.

**Table 1.**
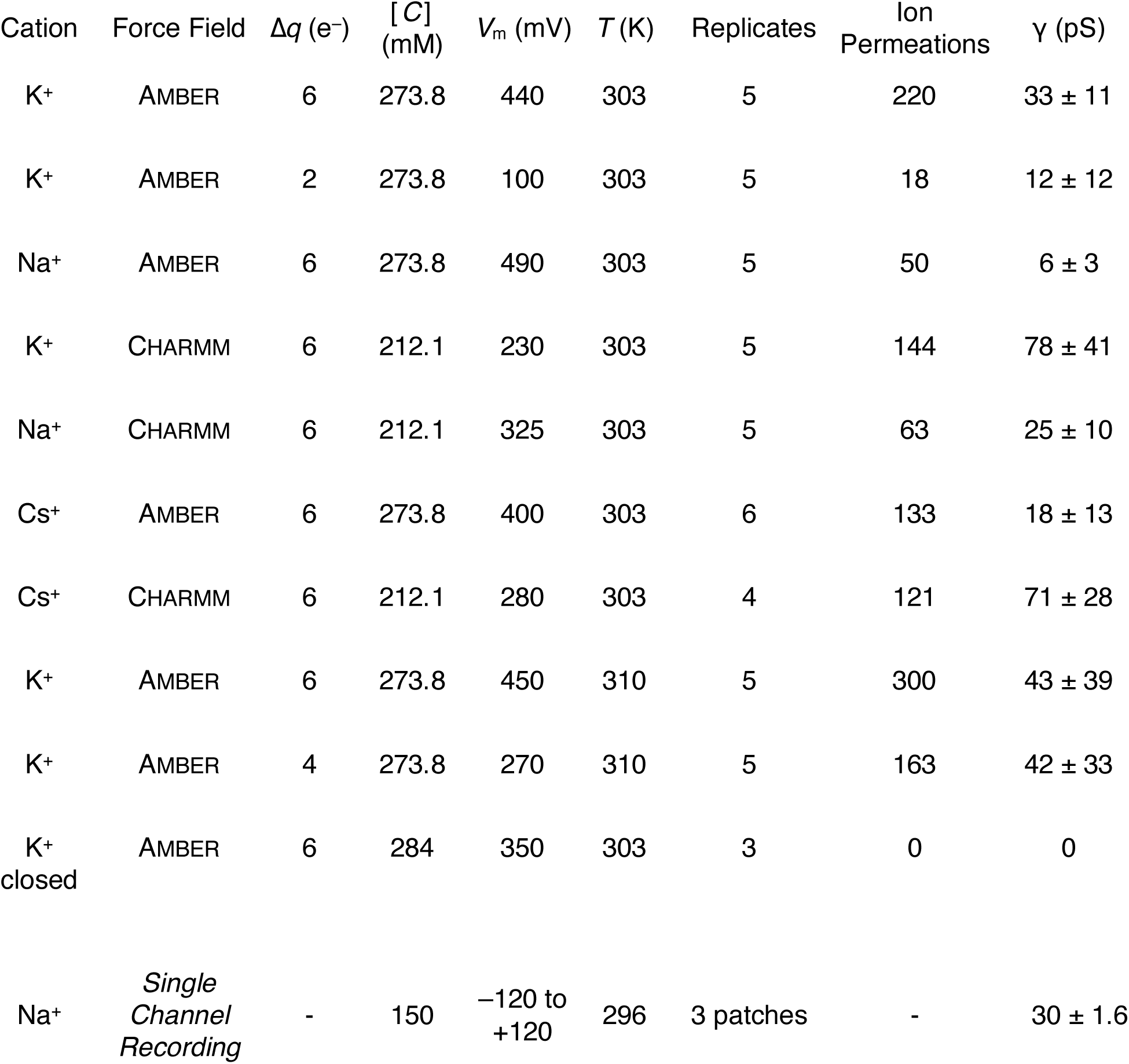
Summary of computational electrophysiology simulations. Simulations of the GluA2 transmembrane domain for individual permeating monovalent cations Na^+^, K^+^ and Cs^+^. In addition to changing cations, we also varied temperature (*T*), charge imbalance (Δ*q*) and the force field employed. *C* is the salt concentration for each simulation. Each simulation replicate was 500 ns except for CHARMM runs where the duration was 250 ns. These simulations represent an aggregate time of 20.5 *µ*s. AMBER refers to AMBER99sb and CHARMM to CHARMM36. For comparison, the conductance of GluA2 (Q) in Na^+^ determined from 3 single channel recordings is also included.

| Cation | Force Field | $\Delta q$ (e<sup>-</sup>) | [C]<br>(mM) | $V_m$ (mV) | $T$ (K) | Replicates | Ion<br>Permeations | $\gamma$ (pS) |
| --- | --- | --- | --- | --- | --- | --- | --- | --- |
| K<sup>+</sup> | AMBER | 6 | 273.8 | 440 | 303 | 5 | 220 | 33 ± 11 |
| K<sup>+</sup> | AMBER | 2 | 273.8 | 100 | 303 | 5 | 18 | 12 ± 12 |
| Na<sup>+</sup> | AMBER | 6 | 273.8 | 490 | 303 | 5 | 50 | 6 ± 3 |
| K<sup>+</sup> | CHARMM | 6 | 212.1 | 230 | 303 | 5 | 144 | 78 ± 41 |
| Na<sup>+</sup> | CHARMM | 6 | 212.1 | 325 | 303 | 5 | 63 | 25 ± 10 |
| Cs<sup>+</sup> | AMBER | 6 | 273.8 | 400 | 303 | 6 | 133 | 18 ± 13 |
| Cs<sup>+</sup> | CHARMM | 6 | 212.1 | 280 | 303 | 4 | 121 | 71 ± 28 |
| K<sup>+</sup> | AMBER | 6 | 273.8 | 450 | 310 | 5 | 300 | 43 ± 39 |
| K<sup>+</sup> | AMBER | 4 | 273.8 | 270 | 310 | 5 | 163 | 42 ± 33 |
| K<sup>+</sup><br>closed | AMBER | 6 | 284 | 350 | 303 | 3 | 0 | 0 |
| Na<sup>+</sup> | *Single*<br>*Channel*<br>*Recording* | - | 150 | -120 to<br>+120 | 296 | 3 patches | - | 30 ± 1.6 |

Although K^+^ permeated readily in the simulations with AMBER99sb force field (20), we observed substantially fewer permeations of Na^+^ under the same simulation condition. The simulated conductances of both K^+^ and Na^+^ were higher in the CHARMM36 force field (21), allowing direct observation of Na^+^ permeation at close to physiological rates (Table 1, Figure 2B). However, the permeation of K^+^ in the CHARMM36 forcefield approaches implausibly high rates, albeit with a very large variability. Higher conductance using CHARMM36 compared to AMBER99sb was previously reported in simulations of K^+^ selective and non-selective NaK-CNG channels (9).

### Binding sites for permeant ions

Selectivity filter architectures differ widely amongst tetrameric ion channels. Whilst K^+^ channels generally have long, straight filters, non-selective cation channels tend to have shorter filters that display more plastic or eccentric geometries (22, 23). To understand how the AMPAR selectivity filter supports cation-selective ion conduction, we determined one- and two-dimensional ion occupancies in the AMPA channel, which were sampled from cumulative simulations of K^+^ and Na^+^ with the respective force fields (Figure 3B, 4A and S2). For each ion species, these plots reveal only one principal binding site (which we call *S*I) in the SF, which is located around 15 Å along the pore axis (Figure 3). At this site, the primary coordination of permeant ions is by backbone carbonyls of Q586 that replace some but not all of waters of hydration (Figure S3). Secondary, less frequent interactions occur with the mobile side chain of Q586 and the backbone carbonyls of Q587. In some trajectories using AMBER99sb force field, solvated Na^+^ ions stay at the *S*_I_ longer than 100 ns (Figure 4B), which may explain the slower permeation. During outward permeation, ions leaving the *S*_I_ site frequently pause above the tip of the SF (25 Å at the pore axis in Figure 4A) and then rapidly enter bulk water as they traverse the upper reaches of the channel. Before reaching the major binding site *S*_I_, multiple ions can simultaneously enter the inner mouth of the region, classically expected to be part of the SF (2, 24), and composed by the residues G588, C589 and D590. K^+^ and Na^+^ paused frequently but briefly at a secondary site (which we denote *S*_II_) which is positioned around 10 Å into the pore, cytoplasmic to the *S*_I_ site (Figure 3B). Compared to the primary site, the *S*_II_ site was weakly populated with shorter residence times (Figure 4B), Therefore we conclude that the true SF region of the AMPA is the shortest seen in any tetrameric channel to date, comprising essentially only Q586 and Q587 for accommodating *S*_I_ and *S*_II_ sites.

**Figure 3.**
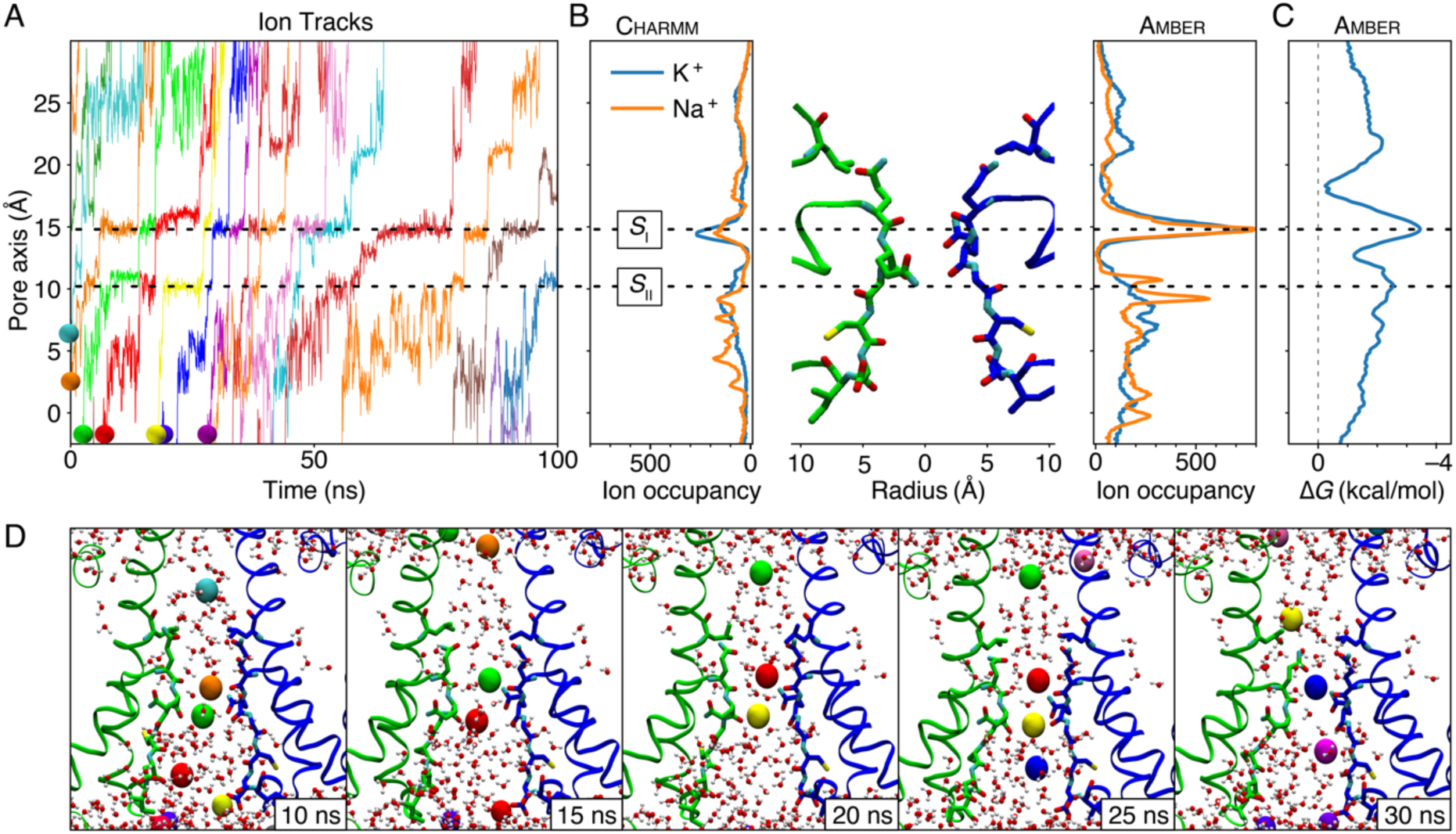
Simulated Na^+^ and K^+^ permeation of GluA2. Simulations were performed at 303 K using an ion imbalance of 6 e^−^ between two compartments *α* and *β*. (A) Representative traces of K^+^ passing through the SF of AMPA channel pore. Ions drawn in panel D are indicated with balls. (B) One-dimensional ion occupancy of cations within the SF of the AMPA channel pore. Left panel: occupancy from simulations with the CHARMM36 force field, right panel from AMBER99sb force field. (C) Free-energy profile for K^+^ permeation constructed from AMBER99sb force field. (D) Representative snapshots of a simulation (from Movie S1) showing multiple K^+^ ions passing the selectivity filter during ion conduction. The color code of ions is identical with the ion track plot in panel A.

**Figure 4.**
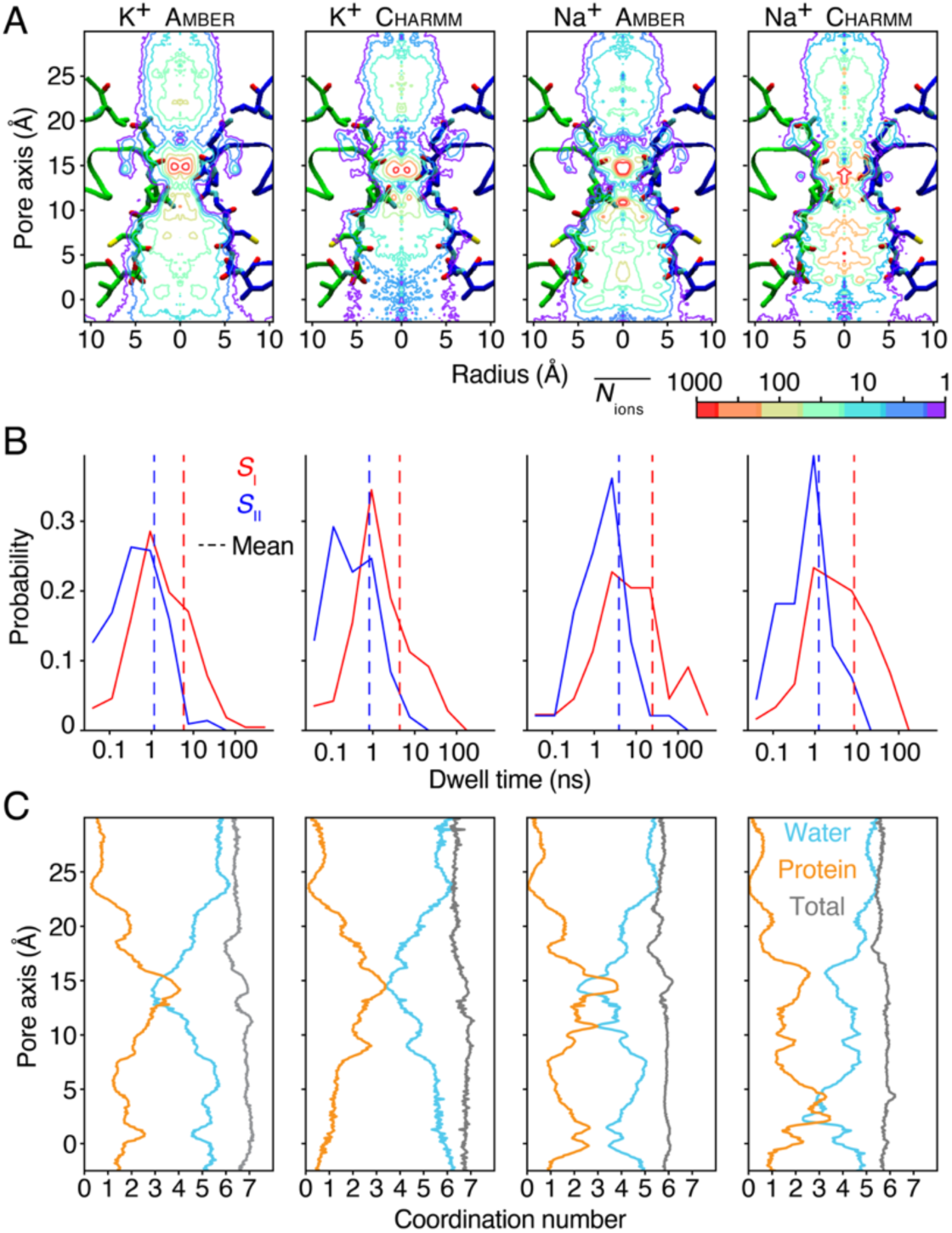
Ion occupancy, dwell time distribution and hydration states. (A) Two-dimensional ion occupancy within the SF resolved radially and in pore axis (z axis) as a contour plot calculated from MD simulations of GluA2 with different ion types (K^+^ and Na^+^) and force fields (AMBER99sb and CHARMM36). The simulations were performed at 303 K using an ion imbalance of 6e^−^ between the two compartments, α and β. The ion occupancy in number of ions per 0.001 Å^3^ per 500 ns was normalized according volume change along radius. (B) Dwell time histograms for *S*I and *S*II binding sites. Vertical dashed lines are means of the respective dwell times. (C) The number of oxygens within the first hydration shell of the according ion type along the pore axis. The number of coordinating water oxygens in blue, of coordinating protein oxygens in orange and the sum in grey.

### Cation conduction mechanism and selectivity

During the ion conduction, most of the time we observed multiple ions reside simultaneously in the canonical SF region of ^586^QQGCDI^591^ (Figure 3D, Movie S1 and S2), with an overall time-average number of 2.4 ± 0.6 and 2.3 ± 0.7 for K^+^ and Na^+^ respectively in the AMBER simulation runs (Figure S4). Ions followed a loosely coupled knock-on mechanism, where in most cases the exit of an ion to the upper cavity was closely followed by the entry of an ion to the SI site (Figure 3D, Movie S1 and S2). From the free-energy profiles (Figure 3C), we conclude that the energy barriers bracketing S_I_ (at around 13 Å and 18 Å along the pore axis) are the major obstacles for ion conduction in the SF. The ion entering the S_I_ site was normally supplied from the adjacent *S*_II_ site, presumably aiding permeant ions to overcome the main energy barriers before and after the *S*_I_ site (6).

To understand better where cation selection occurs, we examined the paths of chloride (Cl^−^) ions during our simulations. Consistent with a very short selective region, Cl^−^ ions occasionally entered the cone-shaped void below the SF and readily approached the QQ-filter from the bundle crossing side. Surprisingly, Cl^−^ ions found stable binding sites immediately above the QQ-filter (Figure S5) in all simulations. We also observed one Cl^−^ ion permeation event (Movie S3), providing a crude estimate of the Cl^−^:K^+^ permeability ratio at around 1:220. In practical terms, this relative selectivity would correspond to an approximately 15 *µ*V shift in the reversal potential (GHK equation, see Methods), compared to no Cl^−^ permeation at all. In other words, at this level of selection between cations and chloride, any reversal potential shifts are too small to be measured experimentally. Further context comes from the observation that R-edited homomeric GluA2 channels are anion permeable (25). The observation of infrequent yet stable anion approach to the face of the QQ-filter favoured for anions by the electric field provides a simple explanation as to how this point mutation can switch selectivity simply by changing local electrostatics.

### Ion hydration states during permeation

To determine the ion hydration states during permeation, we calculated the number of ion-coordinating oxygens from both water molecules and protein residues within each ion’s first solvation shell. Hydration shells are dynamic; here we defined the waters of hydration using the radii corresponding to the minimum in the radius of gyration profiles: 3.1 Å for Na^+^, 3.4 Å for K^+^, and 3.8 Å for Cs^+^ (26, 27). All monovalent ions were principally coordinated by waters in the lower SF region (Figure 4C), while the contribution of protein oxygens (backbone oxygens of Q586, backbone and side-chain oxygens of Q586 and Q587, Figure S3) increases substantially at the major ion binding site and surpasses the water oxygens for K^+^ and Cs^+^. The average number of ion-coordinating oxygens in the first hydration shell (black line) remained remarkably stable along the pore axis, being 6, 7 and 8 for Na^+^, K^+^ and Cs^+^, respectively (Figure 4C). This total number of ion-coordinating oxygens was in very good agreement with the coordination number for hydrated ions determined experimentally (28). The simulations indicate a qualitatively similar mechanism for each ion species. Cations are never fully dehydrated, leading to appreciable water co-permeation. The retention of water is probably a strong factor in determining the lack of selectivity across the alkali metal series.

### Caesium permeation

Classical electrophysiological measurements of reversal potentials for cloned glutamate receptors show at most minor differences in the permeation of Cs^+^ (van der Waals radius 343 pm) from that of Na^+^ and K^+^ (radii 227 pm and 280 pm respectively) (3). We reasoned that to achieve similar permeation statistics despite its greater size, Cs^+^ should either alter selectivity filter geometry, occupy distinct binding sites or have radically different hydration state during permeation. To resolve this point, we simulated Cs^+^ permeation in the same conditions as for K^+^ and Na^+^. From both AMBER and CHARMM simulations we observed a moderately higher Cs^+^ permeation rate (Table 1 and Figure 2B) compared to Na^+^. However, the geometry at the tip of the pore loop did not change. The hydration of Cs^+^ is similar to K^+^, with protein oxygens outnumbering the water oxygens at the *S*_I_ and *S*_II_ sites in the QQ selectivity filter. However, in AMBER simulations the major binding site (*S*_I_) for Cs_+_ is axially displaced by about 0.5 Å towards the bundle crossing compared to those of K^+^ and Na^+^ (Figure 5B, Figure S2), with a concomitant longer distance to the *S*II site. Most critically, in both AMBER and CHARMM simulations, the mean residence time for Cs^+^ at the *S*_I_ site was much less (∼1 ns) than for Na^+^ or K^+^ and matched that at the *S*_II_ site (Figure 4B and Figure 5D). A prediction from its serial binding to two effectively equivalent sites is that Cs^+^ may be selected over Na^+^ and K^+^, and thus be more permeant.

**Figure 5.**
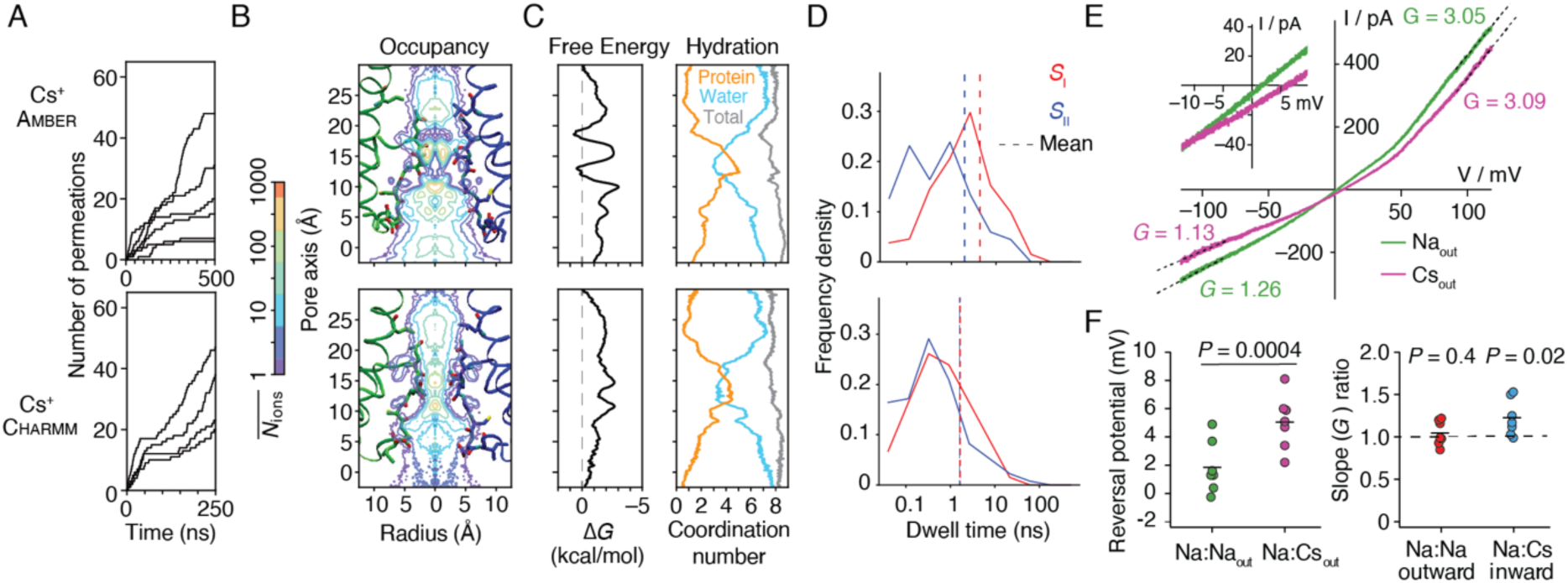
Cs^+^ permeation of GluA2. (A) Permeation events for Cs^+^ in the AMBER99sb and CHARMM36 force fields. (B) Two-dimensional ion occupancy plots (C) Free energy and hydration profiles for Cs^+^ along the pore axis. (D) Dwell time histograms for Cs+ in the *S*I and *S*II sites, with mean dwell time for each site indicated with a dashed line. (E) Outward rectifying responses to voltage ramps in Na^+^ (green) and Cs^+^ (purple) measured with patch clamp electrophysiology. The outward rectification (ratio of chord conductances at +100/–100 mV) was 2.7 ± 0.3 for Na^+^ and 3.1 ± 0.3 for Cs^+^ (*n* = 7 patches). Inset shows the typical shift in reversal potential on exchange of external Cs^+-^ and Na^+^. The pipette solution contained Na^+^. (F) Reversal potential of AMPAR currents in external NaCl and CsCl solution and conductance ratios of Na^+^ versus Cs^+^ for inward and outward permeation. Probability of no difference in the slope ratio was versus a ratio of 1 with paired Student’s t-test.

The permeation of cations in GluA2 (Q) may be different to that observed in extensive studies of GluA1(3). We performed patch clamp electrophysiology experiments on unedited GluA2 channels expressed in HEK 293 cells to reexamine the reversal potentials and slope conductances for Na^+^ and Cs^+^. We readily detected a small but consistent shift in the reversal potential in external Cs^+^ of 3.7 ± 0.5 mV (*n* = 7 patches), meaning Cs^+^ was more permeant than Na^+^ (*P*_Cs_ : *P*_Na_ = 1.16 ± 0.02) like in GluA1. The slope conductance for the inward arm of the I-V relation in Cs^+^ was similar to that of Na^+^ (82 ± 6 %; *n* = 7 patches). These observations are consistent with the AMPAR pore being mildly selective for Cs^+^ over Na^+^ and Cs^+^ retaining a high conductance despite using a subtly different permeation mechanism on the same overall scaffold.

### Conformational flexibility in the vicinity of the SF

Examining the RMSF of heavy atoms in residues in the traditional SF (^585^MQQGCDI^591^) region reveals a clear split in dynamics (Figure 6). The backbone atoms at the pore loop (^585^MQQG^588^) are comparatively immobile, similar to the M3 backbone adjacent at L610. In the lower filter region, the backbone and side chains of CDI residues are comparatively unrestrained. The side chain of Q586 is more flexible than the Q587, as it points toward the cavity and has thus more freedom for movement. Furthermore, the dynamics of the SF were indistinguishable across the different ionic conditions, with the RMSF profiles of both backbone and side-chain atoms in the Na^+^, K^+^ and Cs^+^ simulations being highly similar (Figure 6). These observations further underline that the stable structure of the QQ filter is sufficient for selecting cations.

**Figure 6.**
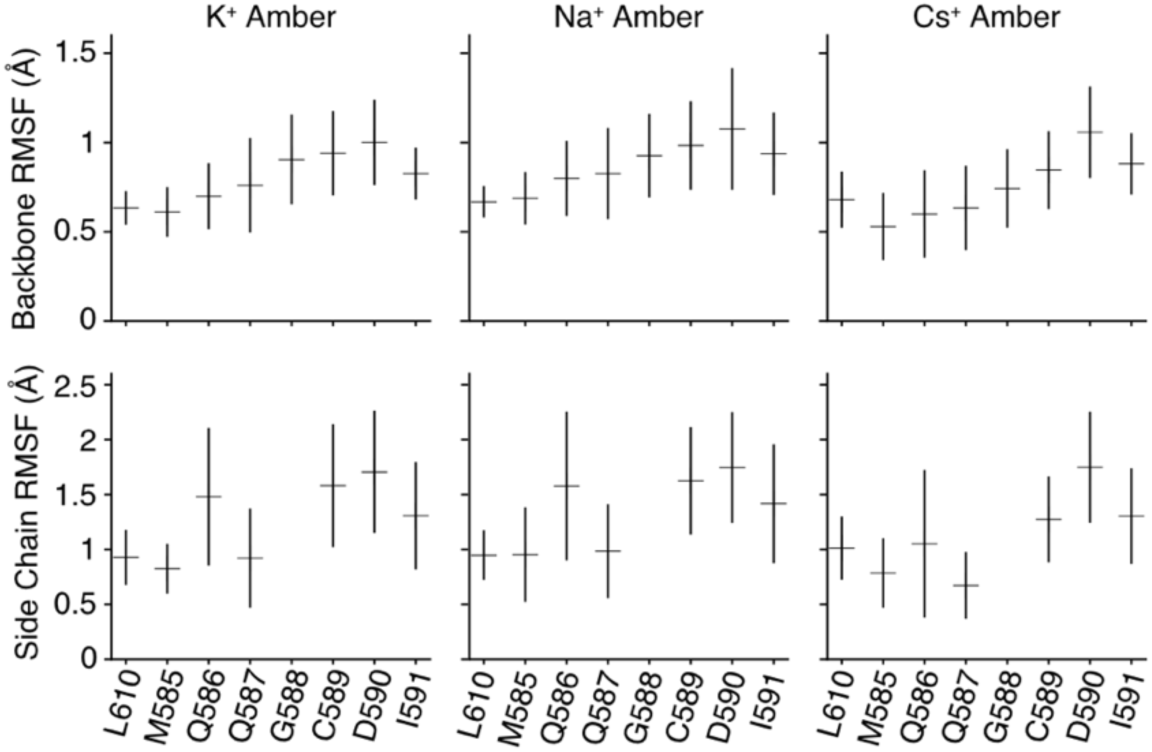
Root mean square fluctuations (RMSFs) of residues surrounding the selectivity filter. RMSFs were calculated separately for backbone and side-chain atoms of selectivity filter residues ^585^MQQGCDI^591^ and L610. The simulations were performed at 303 K using an ion imbalance of 6 e^−^ between the two compartments *α* and *β*.

## Discussion

In this study, we simulated permeation of various monovalent cations through AMPA channel pore using the computational electrophysiology approach. Our direct measurements of ion conduction confirm that the Cryo-EM structure reported by Twomey *et al*. (1) is bona-fide open state, with a conductance large enough to be the fully open state. Particularly, the upper bundle crossing in this structure, which we held open mechanically, provides almost no barrier to conduction. Our simulations provide the first answer to how this channel can allow a range of cations to pass with similar rates. The simulated conductance shows a striking agreement with the single-channel recording data of Na^+^ in GluA2, suggesting that to a first approximation, the ion conduction we observe resembles physiological permeation. The simulations suggested that Cs^+^ permeates with a subtly different mechanism to Na^+^ and K^+^, and we could provide evidence for this using patch clamp electrophysiology.

Although in principle using computational electrophysiology approach we should simulate both inward and outward ion permeations simultaneously from one computational setup, we only observed outward ion flux (from intracellular to extracellular) through the AMPA channel pore in all simulation setups. All simulated conductance listed in Table 1 and Figure 3B are therefore derived from the outward ion permeations. Similar directional flux has also been observed in previous computational electrophysiology simulations of K^+^ selective channels such as KcsA (29). Although electrophysiology reveals that the channel is outward rectifying (Figure 5) for both Na^+^ and Cs^+^, the magnitude of the effect (about 3 fold) is too small to be consistent with an total absence of inward permeation in our simulations. We hypothesize that inward and outward ion permeations may use different SF conformations. In this interpretation, simulations with a longer time scale than performed in the current study would be necessary to reach the conductive states required for inward ion permeations.

For the major physiological ions Na^+^ and K^+^, we observed only one strongly populated ion binding site (*S*_I_) within the SF. This main ion binding site is remarkably congruent with the single ion density observed in the Cryo-EM structure (PDB ID: 5WEO, Figure S6). The centres of the ion densities along the pore axis are within 1 Å. This difference may in part be attributed to the simulations being performed under transmembrane voltages, while the Cryo-EM structure was determined at 0 mV in the absence of ion gradients. In contrast to the major binding site *S*_I_, ion binding to a second on-axis site (*S*_II_) in our simulations was rather short-lived and less ordered. This observation markedly contrasts with the four adjacent ion binding sites of canonical K^+^ channels that were determined by X-ray crystallography (15) and MD simulations (29). In several non-selective cation channels such as NaK (30, 31), CNG (14) and HCN channels (13), the SF consists of either two or three consecutive sites. It has been suggested that a single ion binding site within the NMDA receptor SF is the key to the ion non-selectivity of these cation channels (32). Our finding from the AMPA simulations revealing a single major ion binding site with twin energy barriers is clearly in line with this mechanism. In other words, the AMPA receptor channel is minimally selective and has one of the simplest observed selectivity filters comprising only two residues (Q586 and Q587).

Moreover, we found that during permeation hydration states of the ions in the SF also significantly differ from the ones in the K^+^-selective and several non-selective cation channels. Previous simulations suggested K^+^ pass desolvated through the SF of potassium channels (9), while an entire hydration shell surrounding Na^+^ ions was observed whilst they traverse the SF (9, 27) (10). There is no narrow constriction in the AMPA receptor pore to match those in K^+^ and non-selective NaK channels, with the narrowest portion at Q586 (7.6 Å between CB atoms in 5WEO). Therefore, all monovalent ions remain mostly hydrated in the SF and are only partially dehydrated at Q586. Dehydration of smaller Na^+^ ions requires much more energy (33) (34), and consistent with this expectation, the Na^+^ ions were more hydrated at the QQ filter compared to K^+^ and Cs^+^. The very short SF has a permissive architecture that accommodates the different alkali metals in distinct solvation states.

Based on these observations, we propose that the mechanism of ion conduction through the AMPA channel pore is remarkably different from that seen to date in other tetrameric channels. For K^+^ (29) and several other non-selective cation channels such as NaK (10) and the NaK- CNG mutant (9), ions strictly follow a direct knock-on mechanism when passing through the SF. In contrast, although multiple cations generally occupied the SF-region in the AMPA receptor pore, only one stable axial binding site could be resolved, resulting in a quasi-uncoupled knock-on for K^+^ and Na^+^ conduction. All ions retained a degree of hydration and water was co-permeant. Furthermore, quite distinct from the non-selective NaK channel (10), all monovalent ions investigated in this study (K^+^, Na^+^ and Cs^+^) follow a similar path during permeation, with no requirement for conformational change for physiological permeation rates. The conduction mechanism in the AMPA channel displays some similarities with Na^+^ conduction in the voltage-gated sodium (NaV) channels, where both loosely coupled knock-on and “pass-by” processes compete during Na^+^ permeation (27, 35).

Using extensive MD simulations combined with Markov state model analyses, Furini and colleagues suggested that ion-triggered conformational change within the SF determines ion selectivity in bacterial sodium channels (36). Along the same lines, in our previous study on non-selective NaK channels we proposed that K^+^ and Na^+^ permeation is coupled to distinct SF conformations that are stable in the microsecond time scale and did not readily interconvert (10). In marked contrast to these findings, the short AMPA selectivity filter does not show ion-dependent conformations, and rather, achieves non-selectivity in a quite simple way. The AMPAR provides a minimal selection architecture that is insensitive to ion diameter; the ion hydration shells provide a compensatory variation. However, it is probably not correct to think of the AMPA receptor SF region as a single permissive structure. Instead, the SF of the AMPA receptor is flexible at the nanosecond time scale and allows different ions to permeate using distinct mechanisms but at similar rates. It is also possible that the selectivity filter may be coupled to the upper gate movement or fluctuation as in KcsA (37) and that by holding the upper gate in a single stable conformation, we reduced the SF dynamics. Recent results suggesting that SF perturbations can change the desensitization lifetime are consistent with coupling to regions outside the pore domain (38). On this point, results from CryoEM are inconclusive, with some open and closed structures having similar SF architectures, whereas one open state structure (edited to R586) has marked two fold-symmetry (2). Restraining the linkers was enough to keep the channel stably open, even though we removed flanking Stargazin molecules. Although longer timescales may be required to see any putative relaxation into a TARP-free state, this observation provides further evidence that the modulation of channel gating by Stargazin and other TARPs is principally extracellular (39). Future work will further clarify the relation between conduction properties, receptor conformation and dynamics of the AMPAR pore (40).

## Supporting information

SI Movie 1 Potassium

SI Movie 2 Sodium

SI Movie 3 Chloride

## Methods

### Computational Electrophysiology

To prepare the Cryo-EM structure of the AMPAR in an open state (PDB ID: 5WEO) (1) for MD simulations, we removed the four copies of the auxiliary protein Stargazin. We truncated the receptor linkers before I504 and G774 and after I633 and A820 and added N-methyl amide (NME) and acetyl (ACE) caps at the newly created C- and N-termini. We used MODELLER (41) to build missing loops (residues 550 to 564 of the intracellular loops of each subunits) as well as missing side chains (sidechains of S818 and R819 of subunits B and D, K505 of subunit D). To hold the truncated structure together and in an open state, we restrained protein at the heavy atoms of all four peptide chain termini with a harmonic potential of 10000 kJ/mol.

We performed MD simulations with both CHARMM36 (21) and AMBER99sb (20) force fields. In the AMBER setup, insertion of the AMPAR TMD into a palmitoyloleoyl phosphatidylcholine (POPC) lipid bilayer was performed with the Gromacs internal embedding function, whereas in the CHARMM setup this process was carried out in CHARMM-GUI (42). The concentration of KCl, NaCl and CsCl was 274 mM in simulations with AMBER, except for one simulation setup of the closed channel with the AMBER force field was 284 mM KCl. For CHARMM simulations, we used a salt concentration of 212 mM. Improved lipid parameters (43) and ion parameters (44) were used in the AMBER simulations. We used the Tip3p water model (45) in all simulations.

All the MD simulations were carried out with the GROMACS software package (versions 2016.1 and 2019.5) (46). Short-range electrostatic interactions were calculated with a cutoff of 1.7 nm, whereas long-range electrostatic interactions were treated by the particle mesh Ewald method (47). The cutoff for van-der-Waals interactions were set to 1.7 nm. The simulations were performed at 303 K or 310 K with an enhanced Berendsen thermostat (GROMACS V-RESCALE thermostat) (48). A surface-tension Berendsen barostat was employed to keep the pressure within the membrane-plane (XY-axis) at 250 bar * nm per lipid surfaces and at one bar in z-axis direction (49). All bonds were constrained with the LINCS algorithm (50). Interatomic forces (van-der-Waals and Coloumb) were calculated with a 1.5 nm cutoff. Long range electrostatic forces were computed with the Particle-Mesh-Ewald method. To decrease computational cost, we employed virtual sites for hydrogens in all AMBER simulations. AMBER simulations were calculated with an integration time step of 4 fs. All CHARMM-based simulations used an integration time step of 2 fs.

Before starting the simulation of the AMPAR with an ion gradient, the system was firstly energy-minimized and equilibrated. After the systems energy was minimized to below 1000 kJ/mol with GROMACS “steepest descent” implementation a 10 ns free equilibration without any restraints (except those for the linkers) was performed. For CHARMM-based simulations the recommended energy minimization and equilibration steps from CHARMM-GUI were performed.

For the computational electrophysiology study (17), two copies of the equilibrated system of the AMPAR TMD in the lipid bilayer were included in a simulation box of typically 10 × 10 × 20 nm (Figure 1D). A charge gradient and therefore transmembrane potential was generated by introducing an ion difference of 2, 4 or 6 ions between the two compartments separated by the two lipid bilayers. Since the instantaneous introduction of a charge gradient into the system can produce large perturbations in the system we omitted the first 20 ns of every simulation. During MD simulations, the number of ions in each chamber was kept constant alchemically by an additional algorithm (17). The resulting membrane potential can be calculated by double-integration of the charge distribution using the Poisson equation as implemented in the GROMACS tool *gmx potential* (51). During the simulations a permeation event was counted when an ion traversed the entire filter region. Individual simulation runs were 500 ns in the AMBER setups and 250 ns in the CHARMM setups (Table 1).

The majority of the production runs were conducted at 303 K using a charge imbalance (Δq) of 6 e^−^ (elementary charges) between the two compartments *α* and *β* (Figure 1D). Although ion channel recordings are possible at these voltages, they lay well outside the physiological range. We additionally simulated K^+^ permeation using Δq of 4 and 2 e^−^ with AMBER99sb, respectively, corresponding to a voltage difference of 275 mV and 100 mV. The simulation at 275 mV shows a similar conductance to the one at 450 mV. At 100 mV, the conductance is less but still substantial (Table 1, Figure 2B), indicating that similar conduction mechanisms are at play. The principal consequence of high membrane voltage is merely statistical – it allows us to record more permeation. Furthermore, compared to the simulations conducted at 303 K we noted that simulations at 310 K revealed only slightly larger currents but with much greater variations in the ion permeation rate within a given simulation run (Figure 2B). Finally, to buttress our observations and validate the relevance of the starting structure, we performed equivalent MD simulations of a closed AMPA structure (PDB ID: 5WEM; GluA2-GSG1L-apo-1) with the ATD, LBD and GSG1L removed, and extracellular peptide termini restrained. We observed no ion permeation events over the same time scale as the open structure simulations, and the structure did not spontaneously open on this timescale.

To align results over different setups we used the center of mass of the backbone carbons of the classical selectivity filter residues (^586^QQGCD^590^) as a reference point. All trajectories where analysed with GROMACS tools and PYTHON using MDANALYSIS (52). Molecular visualizations were made with VMD (53).

### Patch Clamp Electrophysiology

HEK293 cells were plated on glass coverslips in dishes and incubated for 20-44 hr before calcium phosphate transfection with 3 *µ*g of cDNA. For macroscopic current recordings, the cDNA transfection was done with the Rat GluA2 (Q) pRK5 vector encoding enhanced green fluorescent protein after an IRES. Single channel recordings from outside-out patches were performed 24 hr after transfection. For single channel recordings, we used a plasmid ratio approach to obtain sparse expression(54), whereby the rat GluA2 Q vector was cotransfected with enhanced green fluorescent protein and empty vector (pCDNA3.1+) in a ratio of 1 : 63 : 313. Standard extracellular solution contained 150 mM NaCl, 0.1 mM MgCl_2_, 0.1 mM CaCl_2_, 5 mM HEPES, 10 *µ*M EDTA and was titrated with NaOH to a pH of 7.3. We included EDTA to chelate trace divalent ion contamination. The internal solution contained 115 mM NaCl, 1 mM MgCl_2_, 0.5 mM CaCl_2_, 10 mM NaF, 5 mM Na_4_BAPTA, 10 mM Na_2_ATP, 5 mM HEPES, titrated to a pH of 7.3 with NaOH. Pipettes were mounted in an ISO holder (G23 Instruments, London, UK) and had a resistance of 3 MΩ for macroscopic current recordings. For single channel recording, pipettes were fire polished to a resistance of 10-25 MΩ and were coated with Sylgard. The junction potential between the pipette and Na+-bath solution (*E*_bath_-*E*_pip_, considering Na^+^, Cl^−^ and F^−^ mobilities) was 3.7 mV (55). For reversal potential experiments with Cs, we substituted NaCl in the extracellular solution with CsCl and titrated the pH to 7.3 with CsOH. CTZ (Hello Bio) was prepared as a 100 mM stock solution in DMSO and used at 100 *µ*M (giving a final concentration of 0.1% DMSO. EDTA stock solution was prepared in NaOH. Reagents were obtained from Carl Roth GmbH (Karlsruhe, Germany), Sigma Aldrich (Munich, Germany), or Toronto Research Chemicals (Toronto, Canada). Ultra-fast perfusion to outside-out patches was used for drug application. The perfusion tools were made with custom designed four-barrel glass (Vitrocom, Mountain Lakes, USA). Currents were filtered at 10 kHz (−3 dB cutoff, 8-pole Bessel) with an Axopatch 200B amplifier (Molecular Devices, Sunnyvale, USA). For analogue-digital conversion an InstruTECH ITC-18 digitizer (HEKA Elektronik Dr. Schulze GmbH, Lambrecht, Germany) was used at 40 kHz sampling rate. Data were recorded and analysed with AxoGraph X (AxoGraph Scientific, Sydney, Australia).

For single channel conductance measurements, in each record, we ran a ramp protocol (–120 to +120 mV, 1.2 V s^−1^) both before and during glutamate application (10 mM) to the outside out patch. The leak current recorded during the no-glutamate ramp was subtracted from the current recorded during glutamate application. Stretches of the recording corresponding to one open channel were selected, the open levels were selected, and fit with a linear relation to obtain the chord conductance.

For macroscopic measurements of reversal potential and conductance, we alternated washing each patch with Cs^+^ and Na^+^ solution. Slope conductances were fit to the traces over the 30 mV ranges at the extreme positive and negative ends of the ramp. For the slope conductance, we compared the conductance ratios (Cs vs Na inward and Na vs Na outward) from the same patch against the null (ratio = 1) using a paired t-test. We assumed that there was a similar junction potential (within 0.5 mV) in both cases, because the junction potential from the pipette to the Cs^+^ solution (–1.7 mV) is cancelled by a second junction potential from Cs^+^ solution back to the bath electrode in Na^+^ (4.9 mV) (56). We added this junction potential to the measured mean reversal potential. The flow rate through the local perfusion tool was low (< 200 *µ*L per minute), meaning the overall bath Cs^+^ concentration remained low.

We used the Goldman-Hodgkin-Katz equation (57) to calculate permeability ratios based on reversal potentials:

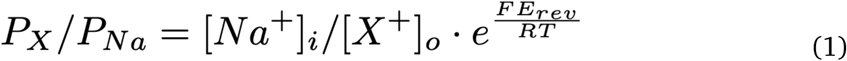

where *X* was Na^+^ or Cs^+^.

Likewise, to calculate the putative shift in *E*_rev_ due to a minor Chloride permeability, we assumed *P*_Cl: Na_ of 1:220 and *P*_Cs: Na_ of 1.16:1, as measured.

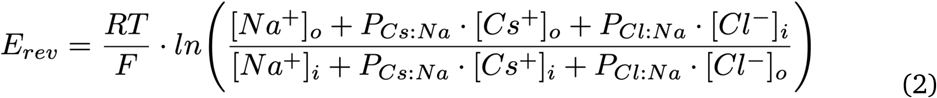

Statistical analysis and data plotting was done in IGOR Pro (Wavemetrics).

## Acknowledgments

This work was funded by the Deutsche Forschungsgemeinschaft (DFG) RU2518 DynIon to A.J.R.P (P3; PL 619/5-1) and H.S. (P3; SU 997/1-1), the DFG under Germany’s Excellence Strategy – EXC 2008 – 390540038 – UniSysCat to H.S, a DFG Heisenberg Professorship (PL 619/3-1) to A.J.R.P. and the ERC CoG “GluActive” (647895) to A.J.R.P. The computations were performed with resources provided by the North-German Supercomputing Alliance (HLRN). The authors gratefully acknowledge the Gauss Centre for Supercomputing e.V. (www.gauss-centre.eu) for funding this project by providing computing time through the John von Neumann Institute for Computing (NIC) on the GCS Supercomputer JUWELS at Jülich Supercomputing Centre (JSC).

The authors declare no competing financial interests.

## Author contributions

All authors designed the experiments, performed the experiments and/or analysed data, wrote and edited the manuscript.

## References

1. Twomey EC, Yelshanskaya MV, Grassucci RA, Frank J, Sobolevsky AI (2017) Channel opening and gating mechanism in AMPA-subtype glutamate receptors. Nature

2. Chen S et al. (2017) Activation and Desensitization Mechanism of AMPA Receptor-TARP Complex by Cryo-EM. Cell 170(6):1234–1246.e14.

3. Jatzke C, Wollmuth LP (2002) Voltage and concentration dependence of Ca2+permeability in recombinant glutamate receptor subtypes. The Journal of Physiology 538(1):25–39.

4. Plested AJR, Vijayan R, Biggin PC, Mayer ML (2008) Molecular basis of kainate receptor modulation by sodium. Neuron 58(5):720–735.

5. Prieto ML, Wollmuth LP (2010) Gating modes in AMPA receptors. J Neurosci 30(12):4449–4459.

6. Bernèche S, Roux B (2001) Energetics of ion conduction through the K+ channel. Nature 414(6859):73–77.

7. Shrivastava IH, Sansom MS (2000) Simulations of ion permeation through a potassium channel: molecular dynamics of KcsA in a phospholipid bilayer. Biophys J 78(2):557–570.

8. Doyle DA et al. (1998) The structure of the potassium channel: molecular basis of K+ conduction and selectivity. Science 280(5360):69–77.

9. Kopec W et al. (2018) Direct knock-on of desolvated ions governs strict ion selectivity in K^+^channels. Nat Chem 10(8):813–820.

10. Shi C et al. (2018) A single NaK channel conformation is not enough for non-selective ion conduction. Nat Commun 9(1):717.

11. Napolitano LMR et al. (2015) A structural, functional, and computational analysis suggests pore flexibility as the base for the poor selectivity of CNG channels. Proc Natl Acad Sci U S A 112(27):E3619–28.

12. Herguedas B et al. (2019) Architecture of the heteromeric GluA1/2 AMPA receptor in complex with the auxiliary subunit TARP γ8. Science 364(6438):eaav9011.

13. Lee CH, MacKinnon R (2017) Structures of the Human HCN1 Hyperpolarization-Activated Channel. Cell 168(1-2):111–120.e11.

14. Li M et al. (2017) Structure of a eukaryotic cyclic-nucleotide-gated channel. Nature 542(7639):60–65.

15. Zhou Y, Morais-Cabral JH, Kaufman A, MacKinnon R (2001) Chemistry of ion coordination and hydration revealed by a K+ channel-Fab complex at 2.0 A resolution. Nature 414(6859):43–48.

16. Saotome K, Singh AK, Yelshanskaya MV, Sobolevsky AI (2016) Crystal structure of the epithelial calcium channel TRPV6. Nature 534(7608):506–511.

17. Kutzner C, Grubmüller H, de Groot BL, Zachariae U (2011) Computational electrophysiology: the molecular dynamics of ion channel permeation and selectivity in atomistic detail. Biophys J 101(4):809–817.

18. Kutzner C et al. (2016) Insights into the function of ion channels by computational electrophysiology simulations. Biochim Biophys Acta 1858(7 Pt B):1741–1752.

19. Verdoorn TA, Burnashev N, Monyer H, Seeburg PH, Sakmann B (1991) Structural determinants of ion flow through recombinant glutamate receptor channels. Science 252(5013):1715–1718.

20. Lindorff-Larsen K et al. (2010) Improved side-chain torsion potentials for the Amber ff99SB protein force field. Proteins 78(8):1950–1958.

21. Huang J, MacKerell AD (2013) CHARMM36 all-atom additive protein force field: validation based on comparison to NMR data. J Comput Chem 34(25):2135–2145.

22. Zubcevic L, Le S, Yang H, Lee SY (2018) Conformational plasticity in the selectivity filter of the TRPV2 ion channel. Nat Struct Mol Biol 25(5):405–415.

23. Liao M, Cao E, Julius D, Cheng Y (2013) Structure of the TRPV1 ion channel determined by electron cryo-microscopy. Nature 504(7478):107–112.

24. Panchenko VA, Glasser CR, Mayer ML (2001) Structural similarities between glutamate receptor channels and K(+) channels examined by scanning mutagenesis. J Gen Physiol 117(4):345–360.

25. Burnashev N, Villarroel A, Sakmann B (1996) Dimensions and ion selectivity of recombinant AMPA and kainate receptor channels and their dependence on Q/R site residues. J Physiol 496 (Pt 1):165–173.

26. Caralampio DZ, Martínez JM, Pappalardo RR, Marcos ES (2017) The hydration structure of the heavy-alkalines Rb^+^ and Cs^+^through molecular dynamics and X-ray absorption spectroscopy: surface clusters and eccentricity. Phys Chem Chem Phys 19(42):28993–29004.

27. Ulmschneider MB et al. (2013) Molecular dynamics of ion transport through the open conformation of a bacterial voltage-gated sodium channel. Proc Natl Acad Sci U S A 110(16):6364–6369.

28. Mähler J, Persson I (2012) A study of the hydration of the alkali metal ions in aqueous solution. Inorganic Chemistry 51(1):425–438.

29. Köpfer DA et al. (2014) Ion permeation in K^+^ channels occurs by direct Coulomb knock-on. Science 346(6207):352–355.

30. Alam A, Jiang Y (2009) High-resolution structure of the open NaK channel. Nat Struct Mol Biol 16(1):30–34.

31. Alam A, Jiang Y (2009) Structural analysis of ion selectivity in the NaK channel. Nat Struct Mol Biol 16(1):35–41.

32. Schneggenburger R (1996) Simultaneous measurement of Ca2+ influx and reversal potentials in recombinant N-methyl-D-aspartate receptor channels. Biophys J 70(5):2165–2174.

33. Biggin PC, Smith GR, Shrivastava I, Choe S, Sansom MS (2001) Potassium and sodium ions in a potassium channel studied by molecular dynamics simulations. Biochim Biophys Acta 1510(1-2):1–9.

34. Kühlbrandt W (2016) Three in a row-how sodium ions cross the channel. EMBO J 35(8):793–795.

35. Boiteux C, Vorobyov I, Allen TW (2014) Ion conduction and conformational flexibility of a bacterial voltage-gated sodium channel. Proc Natl Acad Sci U S A 111(9):3454–3459.

36. Furini S, Domene C (2018) Ion-triggered selectivity in bacterial sodium channels. Proc Natl Acad Sci U S A 115(21):5450–5455.

37. Labro AJ, Cortes DM, Tilegenova C, Cuello LG (2018) Inverted allosteric coupling between activation and inactivation gates in K^+^channels. Proc Natl Acad Sci U S A 115(21):5426–5431.

38. Poulsen MH, Poshtiban A, Klippenstein V, Ghisi V, Plested AJR (2019) Gating modules of the AMPA receptor pore domain revealed by unnatural amino acid mutagenesis. Proc Natl Acad Sci U S A

39. Riva I, Eibl C, Volkmer R, Carbone AL, Plested AJ (2017) Control of AMPA receptor activity by the extracellular loops of auxiliary proteins. Elife 6

40. Kopec W, Rothberg BS, de Groot BL (2019) Molecular mechanism of a potassium channel gating through activation gate-selectivity filter coupling. Nat Commun 10(1):5366.

41. Webb B, Sali A (2016) Comparative Protein Structure Modeling Using MODELLER. Curr Protoc Bioinformatics 54:5.6.1–5.6.37.

42. Jo S, Kim T, Iyer VG, Im W (2008) CHARMM-GUI: a web-based graphical user interface for CHARMM. J Comput Chem 29(11):1859–1865.

43. Berger O, Edholm O, Jähnig F (1997) Molecular dynamics simulations of a fluid bilayer of dipalmitoylphosphatidylcholine at full hydration, constant pressure, and constant temperature. Biophys J 72(5):2002–2013.

44. Joung IS, Cheatham TE (2008) Determination of alkali and halide monovalent ion parameters for use in explicitly solvated biomolecular simulations. J Phys Chem B 112(30):9020–9041.

45. Jorgensen WL, Chandrasekhar J, Madura JD, Impey RW, Klein ML (1983) Comparison of simple potential functions for simulating liquid water. The Journal of Chemical Physics 79(2):926–935.

46. Abraham MJ et al. (2015) GROMACS: High performance molecular simulations through multi-level parallelism from laptops to supercomputers. SoftwareX 1-2:19–25.

47. Darden T, York D, Pedersen L (1993) Particle mesh Ewald: An N· log (N) method for Ewald sums in large systems. The Journal of chemical physics 98(12):10089–10092.

48. Bussi G, Donadio D, Parrinello M (2007) Canonical sampling through velocity rescaling. J Chem Phys 126(1):014101.

49. Zhu Q, Vaughn MW (2005) Surface tension effect on transmembrane channel stability in a model membrane. The Journal of Physical Chemistry B 109(41):19474–19483.

50. Hess B, Bekker H, Berendsen HJC, Fraaije JGEM (1997) LINCS: a linear constraint solver for molecular simulations. Journal of computational chemistry 18(12):1463–1472.

51. Tieleman DP, Berendsen HJC (1996) Molecular dynamics simulations of a fully hydrated dipalmitoylphosphatidylcholine bilayer with different macroscopic boundary conditions and parameters. The Journal of chemical physics 105(11):4871–4880.

52. Gowers R et al. (2016) MDAnalysis: A Python Package for the Rapid Analysis of Molecular Dynamics Simulations. Proceedings of the 15th Python in Science Conference

53. Humphrey W, Dalke A, Schulten K (1996) VMD: visual molecular dynamics. J Mol Graph 14(1):33–8, 27.

54. Groot-Kormelink PJ, Beato M, Finotti C, Harvey RJ, Sivilotti LG (2002) Achieving optimal expression for single channel recording: a plasmid ratio approach to the expression of α1 glycine receptors in HEK293 cells. Journal of Neuroscience Methods 113(2):207–214.

55. Barry PH, Lynch JW (1991) Liquid junction potentials and small cell effects in patch-clamp analysis. J Membr Biol 121(2):101–117.

56. Neher E (1992) [6] Correction for liquid junction potentials in patch clamp experiments. Methods in enzymology 207:123–131.

57. Goldman DE (1943) POTENTIAL, IMPEDANCE, AND RECTIFICATION IN MEMBRANES. J Gen Physiol 27(1):37–60.

