## Supplementary Information for "Non-selective cation permeation in an AMPA-type glutamate receptor"

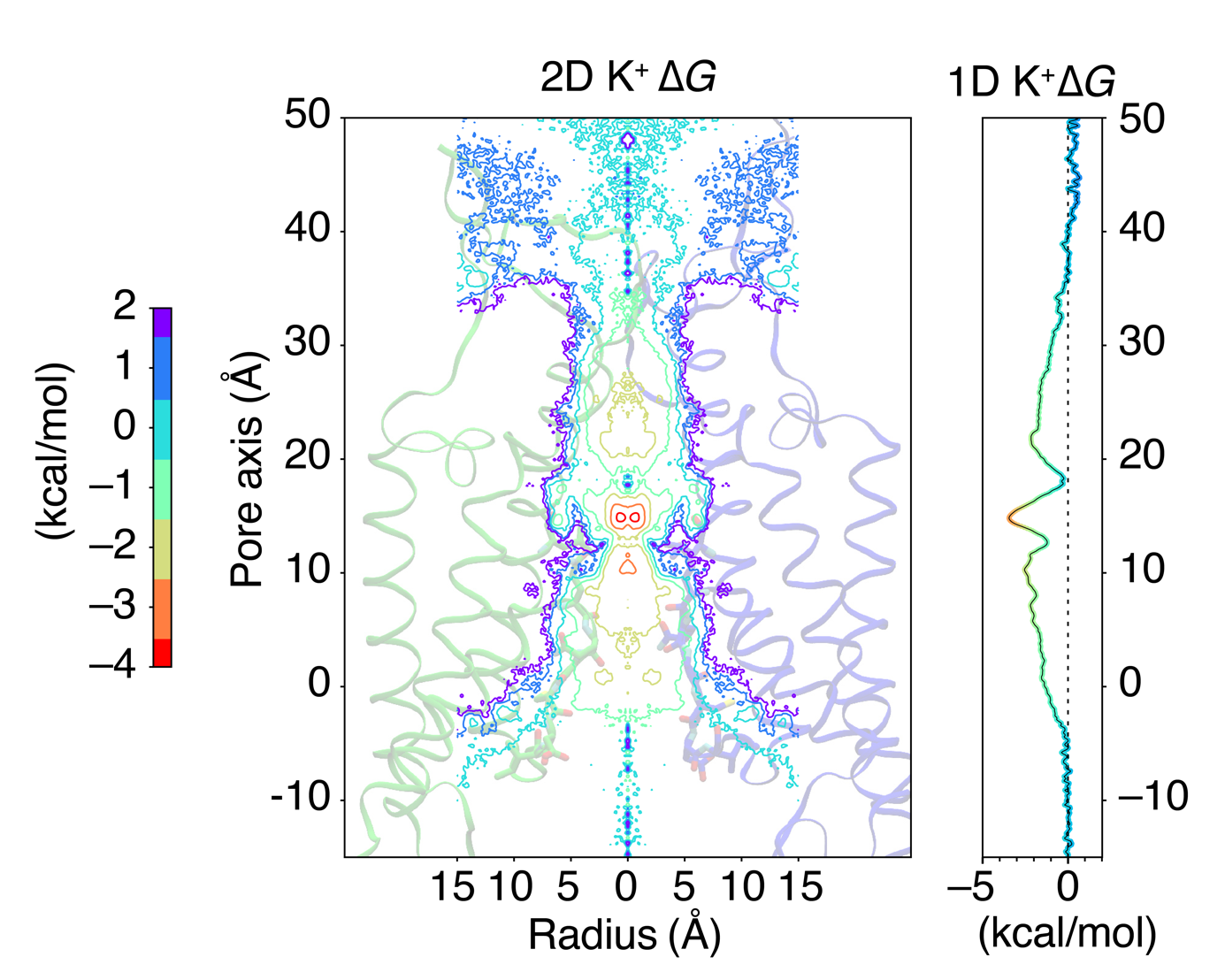

Figure S1. One- and two-dimensional free energy profiles of K^+^ ions within the entire transmembrane region of GluA2, constructed from the total ion positions during the simulations of K^+^ permeation. The binding site at around 14 Å is the only major ion interaction at any point in the pore. The simulations were performed at 303 K, with Amber99sb force field, and using an ion imbalance of 6 e^–^ between two compartments 𝛼 and 𝛽.

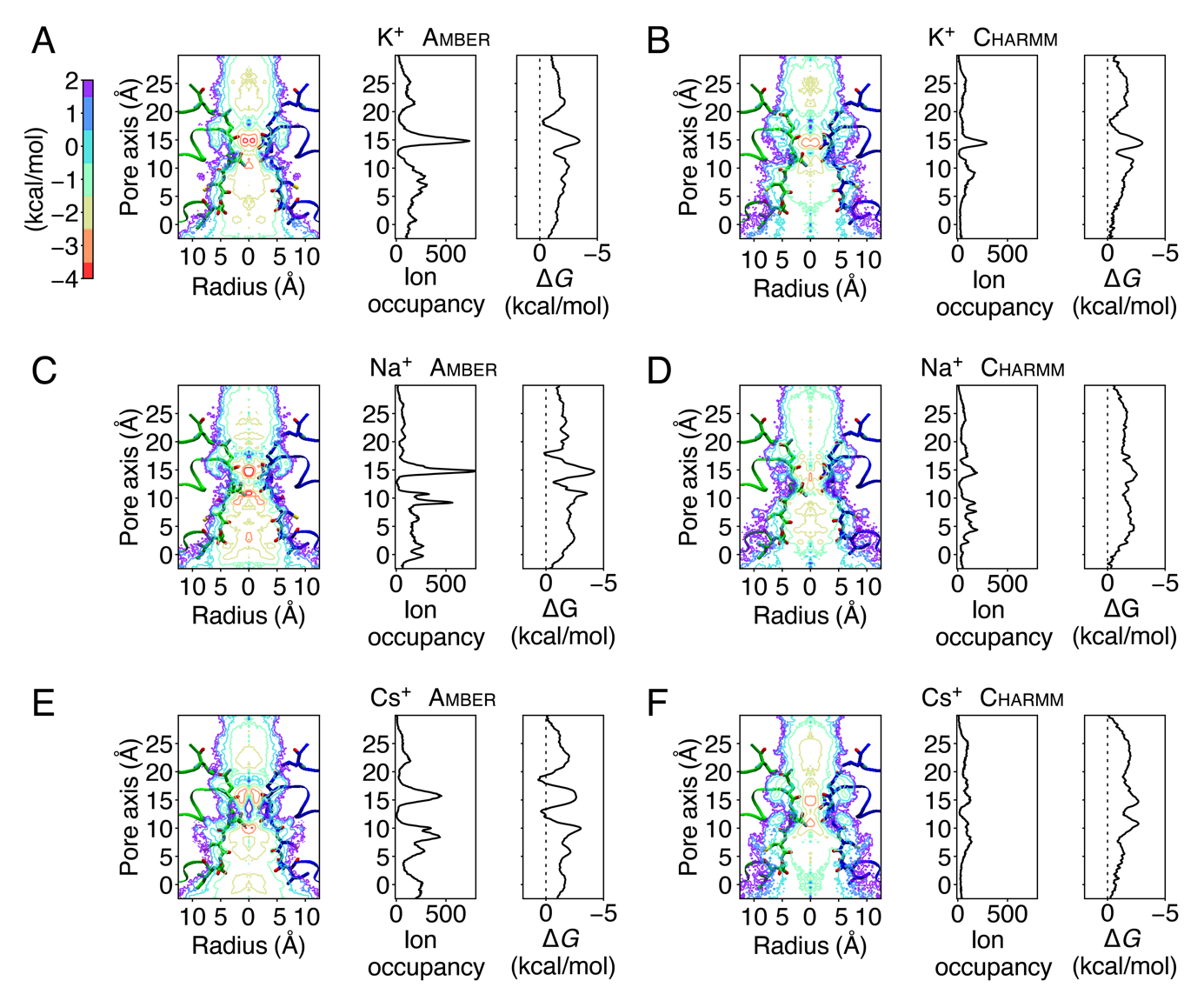

Figure S2. Comparison of two-dimensional projections of ion occupancy with one-dimensional ion occupancy and free-energy profiles of ions within the SF region, constructed from the total ion positions during the simulations. (A) K^+^ permeation using Amber99sb force field, 1D profiles are the same as in Figure 3, 2D projection as in Figure 4. (B) K^+^ permeation using Charmm36force field, 1D profile of ion occupancy as in Figure 3, 2D projection as in Figure 4. (C) Na^+^ permeation using Amber99sb force field, 1D ion occupancy profile is as in Figure 3, 2D projection as in Figure 4. (D) Na^+^ permeation using Charmm36force field, 2D projection as in Figure 4. (E) Cs^+^ permeation using Amber99sb force field, 1D free energy profile and 2D projection as in Figure 5. (F) Cs^+^ permeation using Charmm36 force field, 1D free energy profile and 2D projection as in Figure 5. All simulations were carried out at 303 K using an ion imbalance of 6 e^–^ between two compartments 𝛼 and 𝛽.

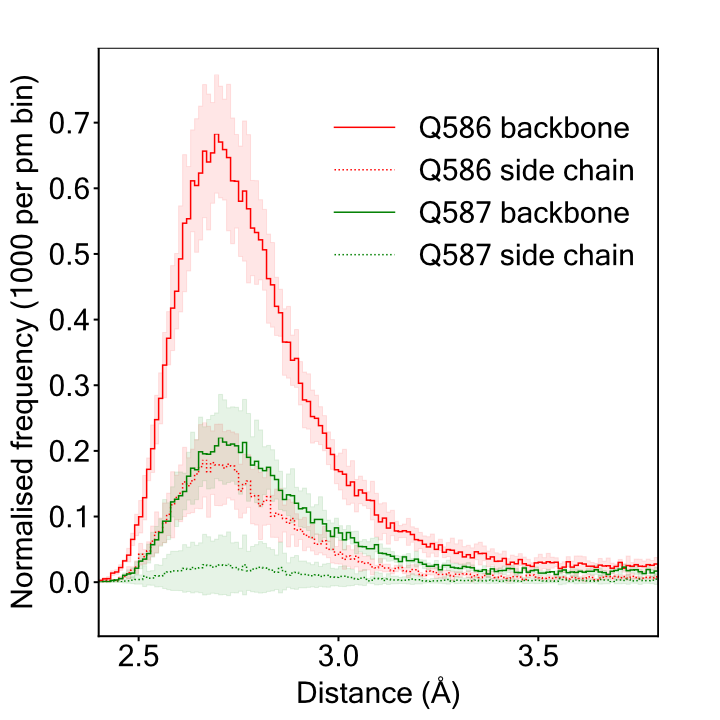

Figure S3. Radial distribution function of distances between permeating K^+^ ions and backbone and side-chain oxygens of Q586 and Q587. The simulations were performed at 303 K, with Amber99sb force field, and using an ion imbalance of 6 e^–^ between two compartments 𝛼 and 𝛽.

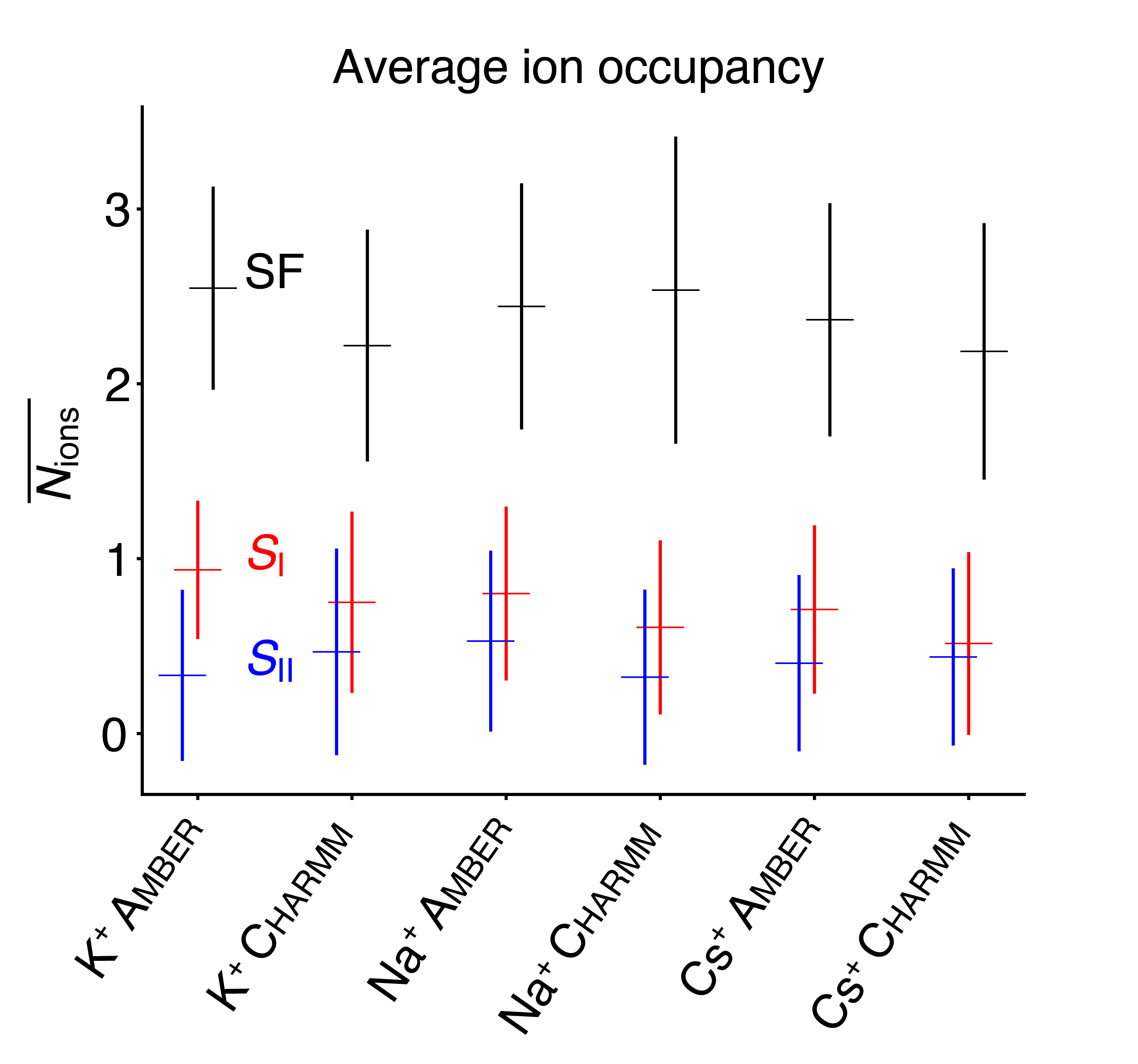

Figure S4. Averaged number of ions in the canonical SF region of the GluA2 pore (^586^QQGCDI^591^) as well as at the ion binding sites *S*_I_ and *S*_II_. All simulations were carried out at 303 K, using an ion imbalance of 6 e^–^ between two compartments 𝛼 and 𝛽, and with Amber99sb and Charmm36 force fields, respectively.

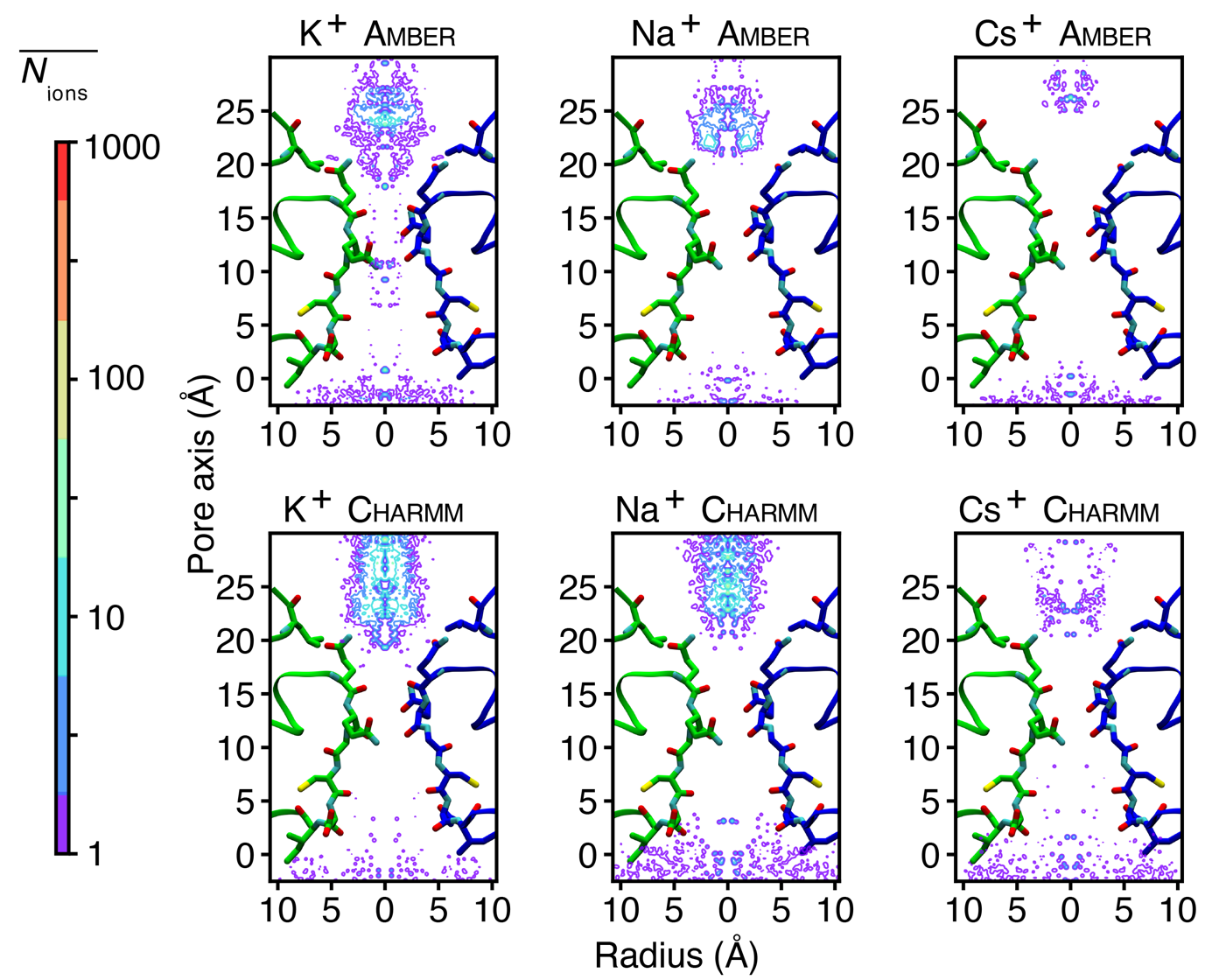

Figure S5. Two-dimensional ion occupancy of Cl^–^ within and around the SF region sampled during K^+^, Na^+^ and Cs^+^ permeation simulations using Amber99sb and Charmm36 force field, respectively. All simulations were carried out at 303 K, using an ion imbalance of 6 e^–^ between two compartments 𝛼 and 𝛽.

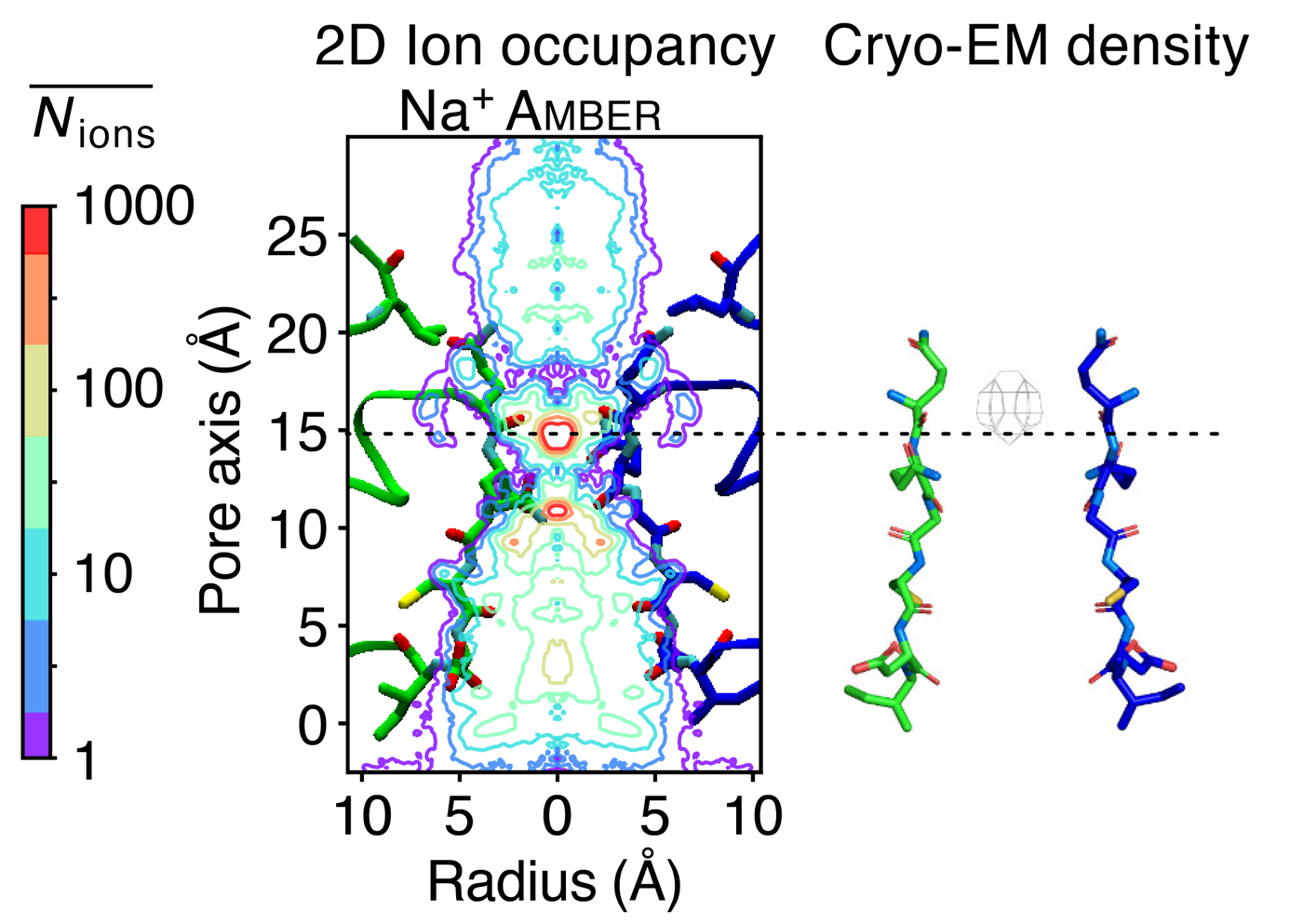

Figure S6. Comparison of the two-dimensional ion occupancy within the SF of GluA2 determined from the Na^+^ simulations using Amber99sb force field and the coulomb density of a presumptive hydrated Na^+^ ion in the Cryo-EM structure (PDB ID: 5WEO). The vertical centre of the main Na^+^ binding site (*S*_I_) is indicated with a dashed line. The simulations were performed at 303 K using an ion imbalance of 6 e^–^ between two compartments 𝛼 and 𝛽.

**Movie Legends**

**Movie S1. K^+^ permeation during a 50 ns trajectory.** Computational electrophysiology simulation using Amber99sb force field under a transmembrane voltage of 350 mV. For clarity, only two subunits of GluA2 are shown in cartoon (blue and green). K^+^ ions are shown in spheres of various colors. Water molecules are drawn as red O bonded to white H atoms.

**Movie S2. Na^+^ permeation during a 50 ns trajectory.** Computational electrophysiology simulation using Charmm36 force field under a transmembrane voltage of 400 mV. For clarity, only two subunits of GluA2 are shown in cartoon (blue and green). Na^+^ are shown in spheres of various colors. Water molecules are drawn as red O bonded to white H atoms.

**Movie S3. Single Cl^–^ permeation observed during a K^+^ permeation trajectory of 50 ns.** The simulation was performed with Amber99sb force field under a transmembrane voltage of 350 mV. For clarity, only two subunits of GluA2 are shown in cartoon (blue and green). K^+^ ions shown as purple spheres and Cl^-^ as green spheres. Water molecules are drawn as red O bonded to white H atoms.
